# Selective dephosphorylation by PP2A-B55 directs the meiosis I - meiosis II transition in oocytes

**DOI:** 10.1101/2020.08.21.260216

**Authors:** S. Zachary Swartz, Hieu T. Nguyen, Brennan C. McEwan, Mark E. Adamo, Iain M. Cheeseman, Arminja N. Kettenbach

## Abstract

Meiosis is a specialized cell cycle that requires sequential changes to the cell division machinery to facilitate changing functions. To define the mechanisms that enable the oocyte-to-embryo transition, we performed time-course proteomics in sea star oocytes from prophase I through the first embryonic cleavage. Although protein levels are broadly stable, dynamic waves of phosphorylation underlie each meiotic stage. We find that the phosphatase PP2A-B55 is reactivated at the Meiosis I/II transition resulting in the preferential dephosphorylation of threonine residues. Selective dephosphorylation is critical for directing the MI / MII transition as altering PP2A-B55 substrate preferences disrupts key cell cycle events after meiosis I. In addition, threonine to serine substitution of a conserved phosphorylation site in the substrate INCENP prevents its relocalization at anaphase I. Thus, through its inherent phospho-threonine preference, PP2A-B55 rewires the cell division apparatus to direct the MI / MII transition.

## Introduction

Animal reproduction requires that oocytes undergo a specialized cell cycle called meiosis, in which two functionally specialized divisions occur in rapid succession to reduce genome ploidy. Following fertilization, the oocyte must then transition to a third and distinct division strategy, mitosis, for early embryonic development. This oocyte-to-embryo transition occurs in a short temporal window, but must achieve high fidelity to ensure that heritable information is accurately transmitted from the parents to the developing embryo. At the center of this progression is a suite of cell cycle regulatory proteins and molecular machines that drive and integrate processes such as chromosome segregation, fertilization, and pronuclear fusion. An important goal is to unravel the complex regulatory mechanisms that precisely coordinate these divisions in time and space within the oocyte. It particular, it remains unknown how phosphorylation and dephosphorylation drive the meiotic divisions allowing oocytes to rewire the cell division machinery at the meiosis I/II transition to facilitate differing requirements.

The female meiotic cell cycle is distinct from mitosis in several ways, necessitating a unique regulatory control. First, oocytes remain in an extended primary arrest in a cell cycle state termed prophase I until receiving an extrinsic hormonal signal (Conti and Chang, 2016; Jaffe and Norris, 2010; Kishimoto, 2018; Von Stetina and Orr-Weaver, 2011). The meiotic divisions use a small, asymmetrically-positioned spindle to partition chromosomes into polar bodies, which do not contribute to the developing embryo (Severson et al., 2016). In addition, the first meiotic division segregates bivalent pairs of homologous chromosomes, whereas for meiosis II, this configuration is reversed and instead sister chromatids are segregated (**Fig. 1A**) (Watanabe, 2012). Finally, meiosis lacks a DNA replication phase between the polar body divisions, which enables the reduction of ploidy to haploid. How the cell division machinery is specialized to perform the distinct functions of meiosis I (MI), and then is rapidly reorganized for the unique requirements of meiosis II (MII) while remaining in meiosis and not exiting into gap or S-phase is an important open question.

**Figure 1.**
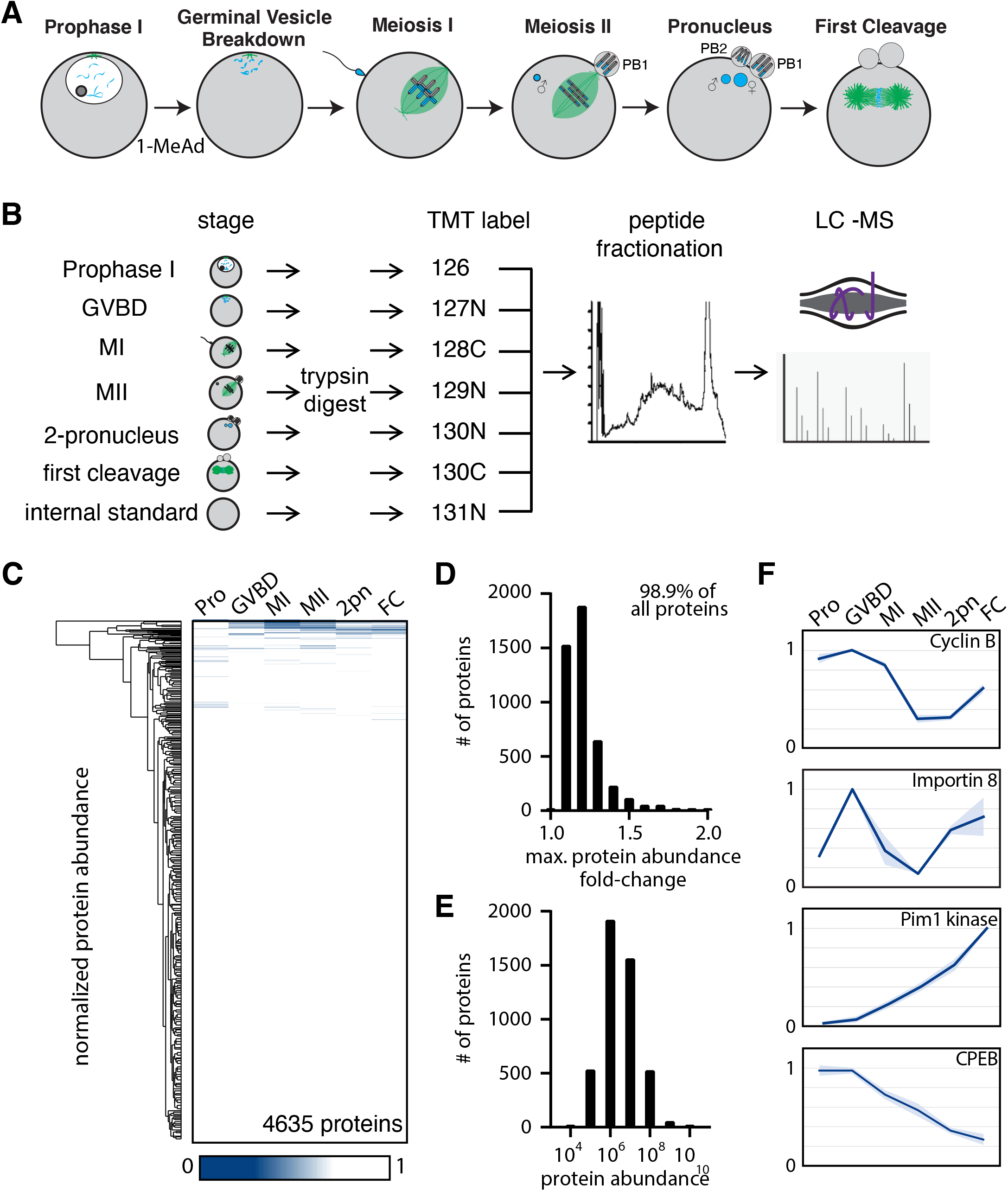
The proteome during the oocyte-to-embryo transition is broadly stable. (A) Schematic of meiotic progression in sea star oocytes, representing the six stages collected for mass spectrometry analysis. (B) Proteomics workflow diagram, in which protein samples were collected in biological triplicate, digested, TMT labelled, fractionated, and analyzed by LC-MS. (C) Hierarchical clustering of the relative abundance of 4,635 proteins detected across three replicates. Individual proteins are clustered (vertically) by the six isolated meiotic stages (horizontally). (D) Histogram of proteins binned by their maximum fold change in abundance, indicating 98.99% of all proteins undergo a fold change of less than 2. (E) Abundance histogram of proteins identified in our analysis reveals a normal distribution. (F) Relative abundances of selected proteins across stages (Pro, Prophase I; GVBD, Germinal Vesicle Breakdown; MI, Meiosis I; MII, Meiosis II; 2-PN, 2 pronucleus; FC, First Cleavage). Light blue shading represents standard deviation.

In this study, we define phosphoregulatory mechanisms that drive the MI/MII transition. We undertook a proteomic and phosphoproteomic strategy using oocytes of the sea star *Patiria miniata*. Prior analyses have revealed proteome-wide changes in animal models including Xenopus, Drosophila, and sea urchins (Guo et al., 2015; Krauchunas et al., 2012; Presler et al., 2017; Zhang et al., 2019). However, the biology of these organisms limits access to a comprehensive series of time points spanning prophase I through the embryonic divisions including the critical meiosis I/II transition. Instead, our sea star proteomics dataset spans the entire developmental window from Prophase I arrest, through both meiotic divisions, fertilization, and the first embryonic division (**Fig. 1A**) (Swartz et al., 2019). We identify a surprising differential behavior between serine and threonine dephosphorylation at the MI/MII transition that we propose underlies key regulatory differences between these meiotic divisions. This regulatory switch is driven by PP2A-B55, which is reactivated after MI to preferentially dephosphorylate threonine residues, thereby creating temporally distinct reversals of Cyclin-dependent kinase and Map kinase phosphorylation. We propose a model in which usage of threonine versus serine endows substrates with different responsivity to a common set of kinases and phosphatases, temporally coordinating individual proteins with meiotic cell cycle progression to achieve specific behaviors for MI and MII without exiting from meiosis.

## Results

### Proteomic analysis reveals stable protein abundance during the oocyte-to-embryo transition

The oocyte-to-embryo transition involves dramatic changes in cellular organization, with an ordered series of events including fertilization, chromosome segregation, polarization, and cortex remodeling. To determine the basis for these cellular changes, we analyzed the proteome composition during the oocyte-to-embryo transition using quantitative tandem mass tag multiplexed mass spectrometry (Thompson et al., 2003). We obtained Prophase I-arrested oocytes from the sea star *Patiria miniata* and treated them with the maturation-inducing substance 1-methyladenine (1-MeAd) to trigger the resumption of meiotic progression in seawater culture (Kanatani et al., 1969). *P. miniata* oocytes progress through meiosis with high synchrony, with key events occurring at stereotypical times following 1-MeAd addition (**Fig. 1A, S1A**). Leveraging these features, we cultured isolated oocytes and collected biological triplicate samples at the following stages: 1) Prophase I arrest (Pro), 2) germinal vesicle breakdown (GVBD), 3) metaphase of meiosis I (MI), 4) prometaphase of meiosis II (MII), 5) just prior to pronuclear fusion (2-PN), and 6) metaphase of the first embryonic cleavage (FC) (**Fig. 1A, S1A**).

We first tested whether protein abundance changes could regulate the oocyte-to-embryo transition (**Fig. 1B**). We identified 8026 total proteins, of which 6212 were identified in two independent time-course series, and 4635 in all three series (**Supp. Table 1, Supp. Fig. 1B**). Surprisingly, we identified few proteins that changed in abundance during these different stages (**Fig. 1C1**). In fact, 98.8% of proteins displayed a maximum fold-change of less than 2 from prophase I to the first embryonic cleavage, with 74.8% of proteins displaying less than a 1.2-fold change (**Fig. 1D**). The absence of changes in protein levels was not due to a bias in our analysis as protein abundance followed a normal distribution (**Fig. 1E**). Despite broad overall stable protein levels, there were several notable exceptions (**Fig. 1F**). For instance, Cyclin B levels were high in Prophase I, GVBD, and MI, but declined sharply in MII, before being partially restored in the first cleavage stage (**Fig. 1F**). These dynamics are consistent with APC/C-mediated destruction of Cyclin B during cell cycle progression (Evans et al., 1983; Okano-Uchida et al., 1998). Gene ontology analysis (Liao et al., 2019) of significantly regulated proteins revealed an enrichment in cytoskeletal proteins, protein involved in RNA binding, and ribosomal components (**Supp. Table 2**). Thus, although some critical regulators of cell cycle progression and other processes fluctuate in their levels, the sea star proteome is broadly stable during the oocyte-to-embryo transition.

### New translation of selected proteins is required for meiotic progression

Although protein levels are largely constant across the oocyte-to-embryo transition, de novo translation could act to maintain steady state levels or may be required to produce a limited set of factors involved in meiotic progression. To test this premise, we globally prevented translation with the 40S ribosomal inhibitor emetine. Emetine-treated oocytes responded to 1-MeAd stimulation to initiate meiosis I, consistent with prior work (Houk and Epel, 1974), but instead of progressing to meiosis II, the maternal DNA decondensed and formed a pronucleus precociously, where they remained arrested even when control oocytes were initiating first cleavage (**Fig. 2A,B**). Based on proteomics of emetine-treated meiosis II or pronuclear stage oocytes, we found that only 108 out of 7,610 proteins identified in our analysis significantly changed with emetine treatment (**Supp. Table 3, Supp. Fig. 2A,B**). These emetine-sensitive proteins fell into diverse categories, but were overrepresented for cytoskeletal elements and actomyosin-related proteins (**Supp. Table 4**). In summary, most proteins are insensitive to translational inhibition, indicating a general lack of turnover between MI and MII, but new protein synthesis is required for progression past meiosis I.

**Figure 2.**
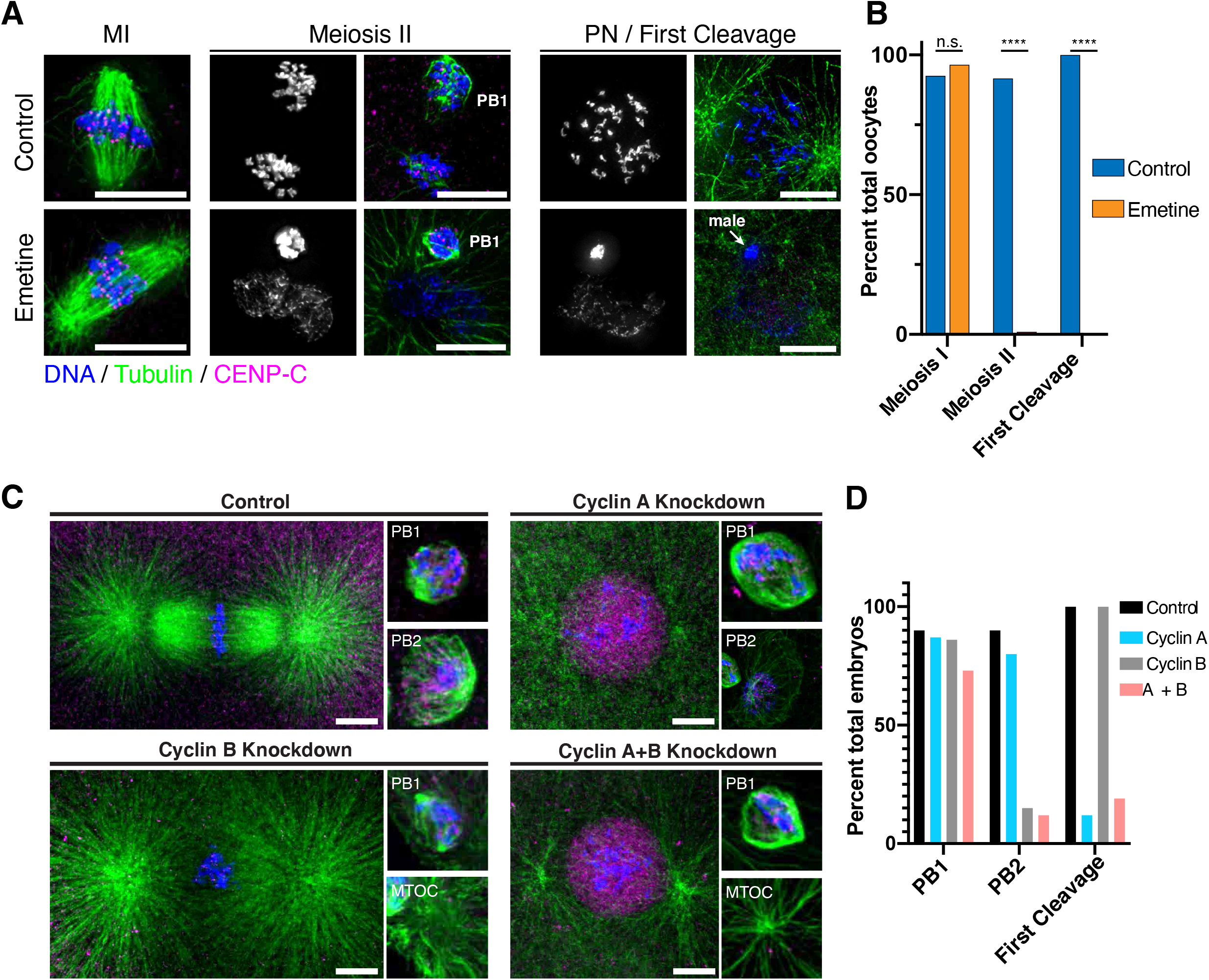
Protein synthesis is required for the MI/MII transition. (A) Immunofluorescence of control or emetine treated oocytes, with DNA provided in single channel grayscale images. While controls proceed to MII and first cleavage, emetine treated oocytes decondense DNA after MI, arrest in a pronuclear state, and fail to incorporate the male DNA. Microtubules were scaled nonlinearly. Scale bars = 10μm. (B) Fraction of oocytes that successfully extruded both polar bodies and underwent first cleavage (Meiosis I control n=107, emetine n=113; Meiosis II control n=107, emetine n=114; First Cleavage control n=31, emetine n=47 oocytes. ****p<0.0001 by Fisher’s exact test). (C) Immunofluorescence of oocytes in which nascent synthesis of Cyclin A, Cyclin B, or both was blocked. Control oocytes extruded both polar bodies and initiated first cleavage. Blocking Cyclin A synthesis did not affect the meiotic divisions, but caused an arrest prior to first cleavage. Blocking Cyclin B instead selectively disrupted the second mitotic division, but the first meiosis and initiation of first cleavage proceeded normally. Combined translational inhibition of both Cyclin A and Cyclin B resulted in an interphase arrest following the first meiotic division. Microtubules were scaled nonlinearly. Scale bars = 10μm. (D) Fraction of oocytes that successfully extrude polar bodies and undergo first cleavage (Cyclin A n=82, Cyclin B n=66, Cyclin A+B n=52 oocytes).

The requirement of protein synthesis for meiotic progression could reflect the need to translate select cell cycle factors. To test whether established cell cycle regulators must be translated de novo, we used morpholino injection to specifically prevent new translation of Cyclin B, one of the few proteins that varies in abundance (**Fig. 2C,D**), as well as Cyclin A, which is synthesized in late MI in a related sea star species (Okano-Uchida et al., 1998). We stimulated oocytes with 1-MeAd immediately following morpholino injection to ensure that pre-existing cyclin protein was unaffected. When new Cyclin A synthesis was blocked, oocytes underwent both meiotic divisions normally and the maternal and paternal pronuclei fused, but these zygotes then arrested with a single, fused pronucleus and failed to progress to the first cleavage (**Fig. 2C,D**). This is consistent with a role for Cyclin A in mitotic entry in cultured cells (Pagano et al., 1992). In contrast, oocytes injected with translation-blocking morpholinos targeting Cyclin B proceeded through MI normally, but failed to extrude the second polar body in MII and thus retained an additional centriole. Surprisingly, these oocytes successfully underwent pronuclear fusion, entered the first cleavage, and formed a mitotic spindle. This suggests that Cyclin B must be translated de novo following Anaphase I to drive meiosis II, but is dispensable for the initial transition from meiosis to embryonic mitosis. Finally, when we simultaneously prevented the new translation of both Cyclin A and B, oocytes completed meiosis I and then arrested in a pronuclear-like state without conducting meiosis II (**Fig. 2C,D**), similar to the effect of translational inhibition by emetine (**Fig. 2A,B**). Taken together, our results suggest that the proteome during the oocyte-to-embryo transition is highly stable, but that the de novo translation of cyclins is required for meiotic progression.

### Defining the phosphorylation landscape of the oocyte-to-embryo transition

The two meiotic divisions, fertilization, pronuclear fusion, and the first mitotic cleavage all occur within less than 3 hours in the absence of substantial changes in protein abundance (**Fig. 1C,D**). This suggests that there are alternative mechanisms to rapidly re-organize the cell division apparatus during these transitions. Therefore, we assessed the role of phosphorylation across meiosis and the first mitotic cleavage (**Fig. 3A**). Our analysis identified a total of 25,228 phosphopeptides across three multiplexed time courses. Among those phosphopeptides, 16,691 were identified in two, and 11,430 in three multiplexes (**Supp. Fig. 1C, Supp. Table 2**). We detected 79.3% of phosphorylation on serine residues, 19.6% on threonine, and 1.1% on tyrosine residues based on a phosphorylation localization probability of ≥0.9 (**Supp. Fig. 1D**). Prior work found similar ratios of S:T:Y phosphorylation based on autoradiographic measurements in chicken cells (92%:7.7%:0.3%) and based on phosphoproteomics in human cells (84.1%:15.5%:0.4%) (Hunter and Sefton, 1980; Sharma et al., 2014).

**Figure 3.**
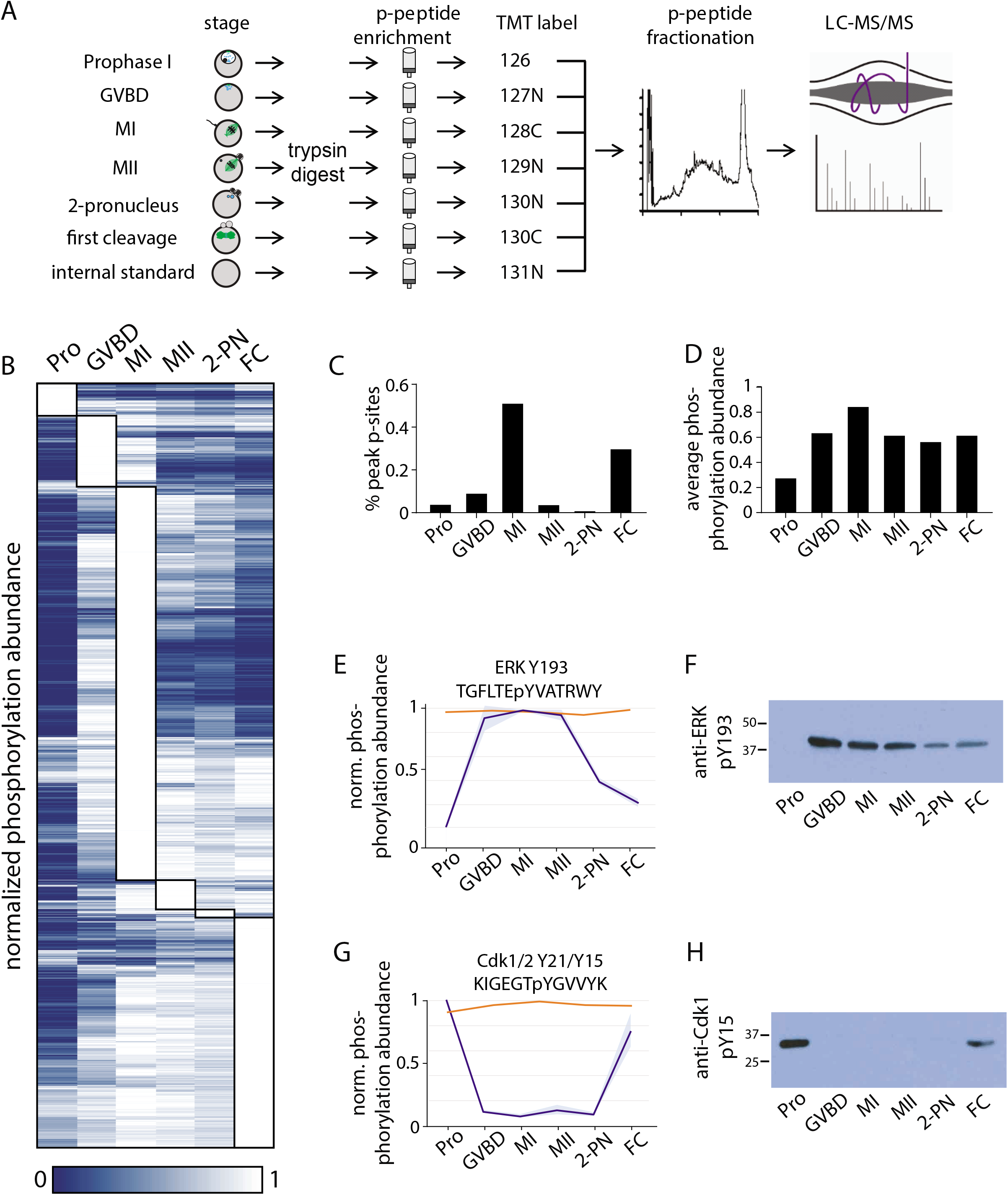
Identification of dynamic phosphorylation changes during the oocyte-to-embryo transition. (A) Proteomics workflow diagram, in which a phosphopeptide enrichment step was performed prior to TMT labelling. (B) Hierarchical clustering of 10,645 phosphorylation events (localization score of 0.9 or higher, p-value of 0.05 or less, and stage specific peak phosphorylation) clustered by phosphosite (vertically) and meiotic stage (columns). (C) Percentage of sites that reach their peak phosphorylation levels across stages, revealing peaks at MI and first cleavage. (D) Average abundance of all phosphorylation events per each stage. (E,G) Temporal phosphorylation levels of conserved sites on the MAP kinase Erk and the kinase Cdk1/2, (blue trace, light blue areas are standard deviation; orange trace represents total relative protein abundance). (H,F) Western blots with antibodies recognizing the inhibitory phosphorylation on Cdk1Y15, and activating phosphorylation on ERK Y193, respectively.

Hierarchical clustering of the dynamic phosphorylation behavior from prophase I to the first embryonic cleavage revealed several striking transitions in global phosphorylation status (**Fig. 3B**). First, Prophase I-arrested oocytes are distinct from the other stages in that they not only display a limited number of phosphorylation sites at the relative maximum phosphorylation levels (**Fig. 3C**), but also have the lowest overall phosphorylation state of the samples tested (**Fig. 3D**). Second, more than half of the total phosphorylation sites identified were maximally phosphorylated in MI, whereas phosphorylation was substantially reduced for MII (**Fig. 3C**). These patterns of phosphorylation imply a critical role for phosphoregulation in specializing the two meiotic and first cleavage divisions, and suggest a role for a low phosphorylation state in maintaining the prophase I arrest.

### Maturation-promoting kinase activity across the oocyte-to-embryo transition

To remain arrested in Prophase I, Cyclin-dependent kinase 1 (Cdk1) and maturation promoting kinases must kept inactive. As our proteome analysis indicated that the majority of proteins in the oocytes, including kinases, are present constitutively (**Fig. 3B,E,G**), kinase activity must be controlled post-translationally. Therefore, we next analyzed the pattern of phosphorylation events on established cell cycle kinases. The meiotic divisions and secondary arrest that occurs in the absence of fertilization require MAP kinase activity downstream of the conserved activator Mos (Dupré et al., 2011; Tachibana et al., 2000). Based on our phosphoproteomics and Western blotting, we found that a conserved activating phosphorylation on the MAP kinase p42/ERK phosphorylation (Y204) was undetectable in Prophase I-arrested oocytes, but high in meiosis I and II (**Fig. 3E,F**). Although phosphorylation is low globally in Prophase I-arrested oocytes, we found high phosphorylation for inhibitory sites on Cdk1 or Cdk2 (Y21 or Y15, respectively) based on our phosphoproteomic analysis and Western blotting using phospho-specific antibodies against these conserved sites (**Fig. 3G, H**). The pattern of these phosphorylation events suggests that p42/ERK and Cdk are inactive in Prophase I-arrested oocytes, but are high throughout the meiotic divisions. Finally, meiotic resumption in sea star oocytes requires Greatwall kinase (Kishimoto, 2018). Greatwall is sequestered in the germinal vesicle in Prophase I sea star oocytes and is activated downstream of Cdk1/Cyclin B (Hara et al., 2012). We identified a conserved activating phosphorylation within the activation segment of Greatwall kinase (T194 in humans; T204 in sea star) (**Supp. Fig. 3A**) (Blake-Hodek et al., 2012; Gharbi-Ayachi et al., 2010), indicative of high Greatwall kinase activity during GVBD and MI, and reduced activity in later stages. In summary, our phosphoproteomic time-course reveals orchestrated transitions in the activity of regulatory kinases during the oocyte-to-embryo transition.

### Prophase I arrest is enforced by high phosphatase activity

Our phosphoproteomics analysis indicated that Prophase I is characterized by low global phosphorylation. To determine whether this state is reinforced by phosphatase activity to maintain the primary arrest, we analyzed our dataset for modifications to the major cell cycle phosphatases, PP1 and PP2A (Nasa and Kettenbach, 2018), and their regulators. PP1 activity is inhibited by phosphorylation of both its catalytic subunit (phosphorylation of T320 (human) by Cdk1 (Dohadwala et al., 1994; Kwon et al., 1997); **Supp. Fig. 3B**) and of its regulatory subunit NIPP1 (**Supp. Fig. 3C-E**). PP1 T316 (corresponds to T320 in humans) is notably hypo-phosphorylated in Prophase I-arrested oocytes (**Supp. Fig. 3B;** (Beullens et al., 1999; O’Connell et al., 2012)), which would result in high PP1 activity. Similarly, NIPP1 phosphorylation is low in Prophase I-arrested oocytes (**Supp. Fig. 3C,D**). Preventing NIPP1 phosphorylation with phospho-inhibitory mutations (*P. miniata* NIPP1 S199A or S197A S199A double mutants) resulted in a >5-fold increase in their binding to human PP1 when expressed in human 293T cells (**Supp. Fig. 3F**). Thus, the low NIPP1 phosphorylation in Prophase I oocytes would increase the association between PP1 and NIPP1 to modulate its phosphatase activity towards specific substrates.

We also identified phosphorylation events predicted to inhibit PP2A activity through its B55 subunit (**Supp. Fig. 3G**). Upon phosphorylation of ARPP19 by Greatwall kinase, phospho-ARPP19 (S67 in human, S106 in *P. miniata)* binds to and inactivates the regulatory PP2A regulatory subunit B55 (Gharbi-Ayachi et al., 2010; Okumura et al., 2014). Indeed, we found that PmARPP19 thio-phosphorylated *in vitro* by Greatwall kinase (Gharbi-Ayachi et al., 2010) resulted in increased Cdc25 phosphorylation when added to mitotic human cell lysates (**Supp Fig. 3H**). Thus, phospho-ARPP19 S106 acts an active B55 inhibitor such that ARPP19 phosphorylation can serve as a proxy for assessing PP2A-B55 activity in sea star oocytes. *Pm*ARPP19 S106 phosphorylation is low in Prophase I-arrested oocytes, but is high in GVBD and MI, before decreasing again in MII (**Supp. Fig. 3I**). Therefore, we conclude that PP2A-B55 activity is high in prophase I, low for MI, but reactivated following the MI/MII transition.

To test the functional requirement for these phosphatases in maintaining prophase I arrest, we treated oocytes with the potent PP1 and PP2A inhibitor Calyculin A. Following Calyculin A addition, we found that 100% of oocytes spontaneously underwent GVBD within 70 minutes (**Supp. Fig. 4A**; also see (Tosuji et al., 1991)). However, the oocytes failed to progress past this GVBD-like state based on the absence of a contractile actin network, chromosome congression, or spindle formation (**Supp. Fig. 4B**). Therefore, while phosphatase activity is required to maintain a normal prophase I arrest, its inhibition is not sufficient to recapitulate physiological meiotic resumption. Calyculin A treatment resulted in the broad upregulation of phosphorylation based on a phosphoproteomics analysis (**Supp. Table 5**), with 85% of phosphorylation sites being maximally phosphorylated after 70 min following Calyculin A treatment (**Supp. Fig. 4C**). Indeed, the phosphorylation events associated with the different meiotic stages increased, whereas those associated with the Prophase I arrest instead decreased (**Supp. Fig. 4D**). Collectively, with phosphoproteomic and functional analyses indicates that there is high activity of PP1 and PP2A-B55 in Prophase I-arrested oocytes that enables the low global phosphorylation state, with additional waves of phosphatase activity controlling subsequent phases of meiotic phosphorylation.

### PP2A-B55 drives selective dephosphorylation at the MI/MII transition

During meiosis, oocytes must undergo two consecutive chromosome segregation events without exiting into an interphase state. These rapid divisions occur within 30 minutes of each other in the sea star, but they each achieve distinct functions. Therefore, some MI-associated phosphorylations must be reversed to allow progression to MII, while others must be maintained to remain in meiosis. Of the stages tested, MI displayed the largest number of sites with maximal phosphorylation (**Fig. 3A, 4A**), including a large number of sites with a TP or SP consensus motif indicative of CDK or MAPK-dependent phosphorylation. Hierarchical clustering of sites maximally phosphorylated in MI revealed three distinct clusters. Strikingly, a subset of sites sharply decreased in phosphorylation after MI (**Fig. 4B**, Cluster 2), whereas other sites remain phosphorylated during MII and the first embryonic cleavage (**Fig. 4B**, Cluster 3). A third cluster displayed intermediate dephosphorylation kinetics (**Fig. 4B**, Cluster 1). These differential behaviors could reflect a mechanism by which the meiotic divisions are specified and underlie the transition from MI to MII.

**Figure 4.**
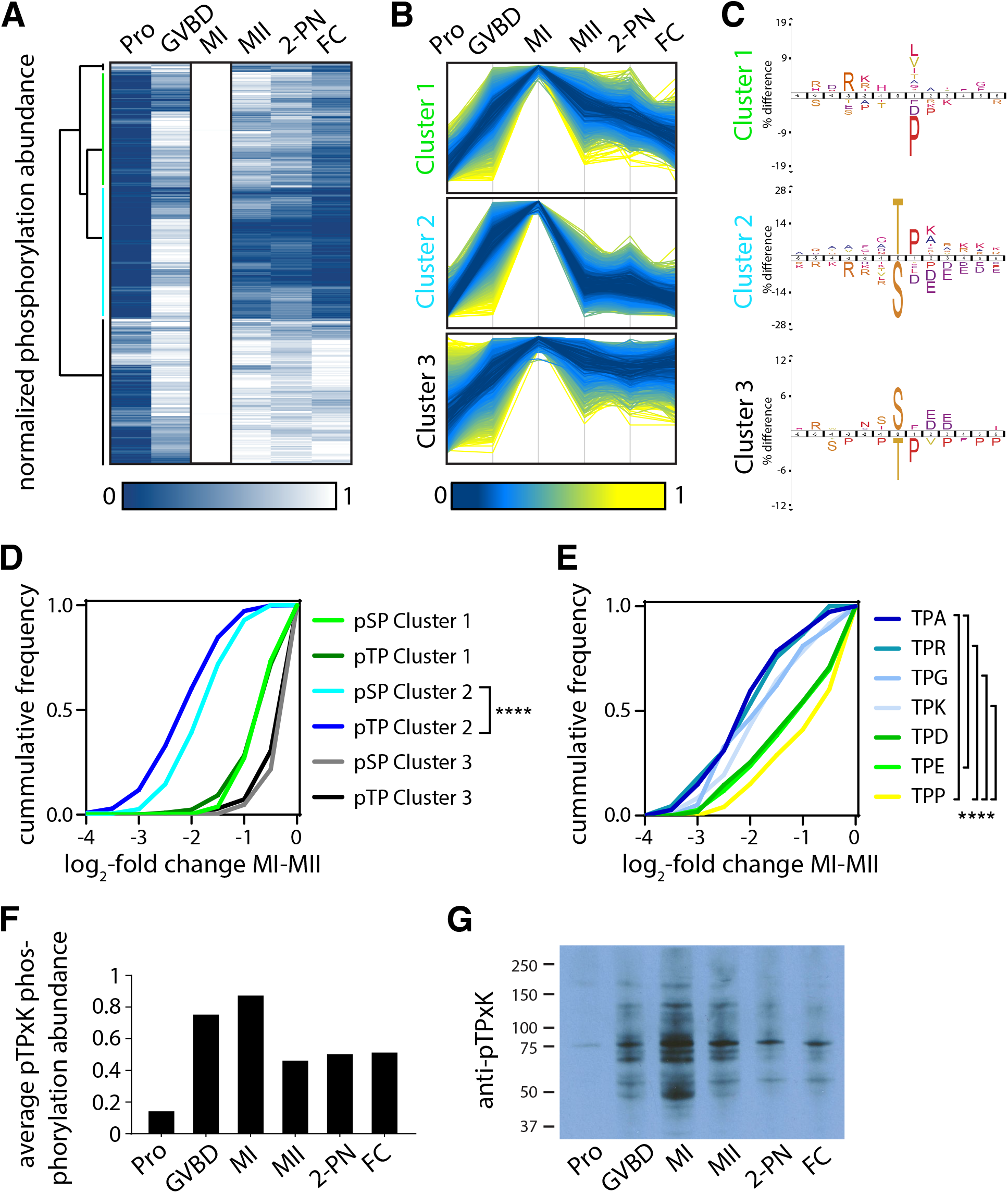
Serine and threonine display distinct phosphorylation behaviors. (A) Heatmap representation of a subset of sites that peak in phosphorylation in meiosis I. Hierarchical clustering identifies three phosphorylation clusters with distinct temporal behaviors, indicated by green (cluster 1), cyan (cluster 2), and black (cluster 3) vertical lines. (B) Line graphs of temporal phosphorylation levels of sites within each of the three clusters. Color scale represents the distance from the mean. (C) Sequence logos for over and underrepresented motifs within the three clusters. Threonine with proline in the +1 position followed by basic amino acids is overrepresented in Cluster 2, which is dephosphorylated after MI. In contrast, Cluster 2 is depleted for serine followed by acidic amino acids. Instead, Cluster 3, which is more stably phosphorylated, is enriched for serine but depleted for threonine as the phosphoacceptor. (D) Cumulative frequency distribution of phosphopeptides with proline-directed serine or threonine phosphorylation. Significant differences in the population distribution as determined by KS statistics (p-value: **** <0.0001). Only cluster 2 shows a significant difference in the dephosphorylation of SP versus TP phosphorylation sites. (E) Cumulative frequency distribution of phosphopeptides with proline-directed threonine phosphorylation with different amino acids in the +2 position. Only the most significant differences are indicated (p-value: **** <0.0001). (F) The average phosphorylation abundance of sites that peak in MI, comparing threonine (black bar) vs. serine (gray bars) phosphorylation sites across stages as determined by phosphoproteomics. (G) Western blot with an antibody recognizing phosphorylated threonine-proline followed by lysine in the +3 position, revealing multiple bands that peak in MI.

To determine the mechanisms controlling these behaviors, we analyzed the specific phosphorylation events that are eliminated after MI (**Fig. 4B**, Cluster 2) compared to those that are retained (**Fig. 4B**, Cluster 3). Motif analysis of Cluster 2 sites revealed that they predominantly occur on threonine with proline in the +1 position (TP sites). Furthermore, we identified an enrichment for a basic amino acids starting in the +2 position (e.g. lysine and arginine) and a depletion of acidic amino acids (e.g. aspartic and glutamic acid) (**Fig. 4C**). This phosphorylation site preference is consistent with the consensus motif dephosphorylated by PP2A-B55 that we and others recently reported (Cundell et al., 2016; Kruse et al., 2020; McCloy et al., 2015). In contrast, MI-specific phosphorylation sites are depleted for serine and for downstream acidic residues. We next directly compared the phosphorylation of SP versus TP sites in the three clusters and found that TP phosphorylation declines more substantially than SP after MI (**Fig. 4D**). Notably, we also observed that the nature of the amino acids in the +2 position of TP site determined its dephosphorylation. TP sites with a small non-polar or basic amino acid were significantly more dephosphorylated than TP with acidic amino acids or proline (**Fig. 4E, Supp. Figure 5A**). On average, the mean phosphorylation of TPxK motifs detected by phosphoproteomics peaked in MI, but then declined for MII (**Fig. 4F**). In further support of this analysis, Western blots using antibodies against phosphorylated pTPxK sites indicated a peak in MI, followed by strong reduction in threonine phosphorylation in MII (**Fig 4G**). Taken together, these data reveal unexpected behavioral differences for SP vs. TP phosphorylation sites at the MI to MII transition, with threonine residues followed by basic or nonpolar amino acids being preferentially dephosphorylated. These dephosphorylation behaviors are consistent with regulation by PP2A-B55, suggesting that this phosphatase must be reactivated at the MI/MII transition.

PP2A-B55 inhibition is required for meiotic resumption from Prophase I into MI in response to hormonal stimulation (Hara et al., 2012; Okumura et al., 2014), but a role for PP2A-B55 at the MI/MII transition has not been defined. To determine the activation state of PP2A-B55 at the MI/MII transition, we assessed its conserved regulatory pathway. Cdk1 phosphorylation activates Greatwall kinase, which then phosphorylates and activates the B55 inhibitor ARPP19 (**Fig. 5A, Supp. Fig. 3A**) (Hara et al., 2012; Okumura et al., 2014). Therefore, a drop in Cyclin B levels and CDK activity would drive the reactivation of PP2A-B55. Indeed, following MI, our dataset reveals a decrease in Cyclin B levels, and a corresponding reduction in the activating phosphorylations on Greatwall kinase (T204) and ARPP19 (S106). This would result in the release of PP2A-B55 to dephosphorylate its substrates (**Fig. 5B, 1F, Supp. Fig. 3A,I**). Together, these observations suggest that PP2A-B55 is reactivated at the MI/MII transition through a conserved pathway involving ARPP19, Greatwall, and CDK/Cyclin B (**Fig. 5B**).

**Figure 5.**
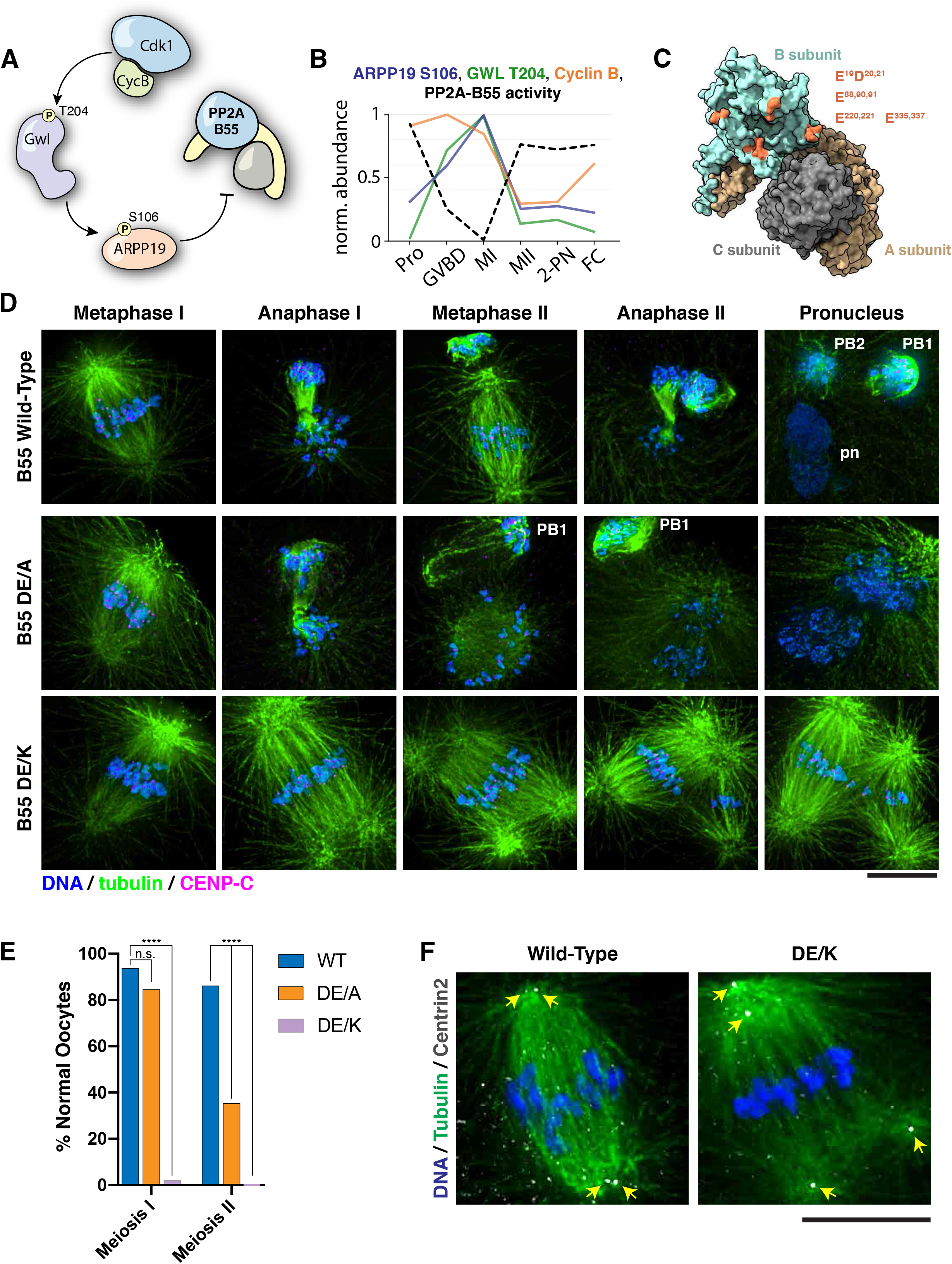
PP2A-B55 substrate specificity is required for the MI/MII transition. (A) Schematic of the Gwl-ARPP19 pathway regulating PP2A-B55. (B) Relative phosphorylation levels of ARPP19 S106, Gwl T204, and protein abundance of Cyclin B. Individual traces with standard deviation are in Supp. Fig S5A,B. PP2A-B55 activity trace values are derived from the substrates in Supp. Fig. 6B-D (1-mean phosphorylation levels). (C) Crystal structure of PP2A holoenzyme with B subunit colored X. Mutated residues within the acidic surface are indicated. (D) Immunofluorescence of meiotic time-course of oocytes expressing wild-type or mutant B55 constructs through. In contrast to wild-type oocytes, DE/A mutants do XYZ. The DE/K mutant oocytes successfully form the first meiotic spindle, but fail to undergo homologous chromosome or sister centromere separation. The spindle poles do separate, ultimately resulting in a two semi-distinct spindles. Microtubules were scaled nonlinearly. (E) Percentage of oocytes successfully completing MI and MII (control n=65, DE/A n=65, DE/K n=49 oocytes; ****p<0.0001 by Fisher’s Exact Test). (F) Centrin2 staining of B55 wild-type and DE/K mutant expressing oocytes stained for Centrin 2 reveals centriole separation at the spindle poles. Scale bars = 10μm.

### Selective dephosphorylation by PP2A-B55 is required for the MI/MII transition

Our data are consistent with a model in which PP2A-B55 reactivation at the MI / MII transition drives the selective dephosphorylation of substrates to rewire the cell division machinery and distinguish these divisions. We therefore sought to test the functional contribution of PP2A-B55 to the MI / MII transition. As small molecule inhibition of PP2A also inhibits other phosphoprotein phosphatases and resulted in a failure in meiotic progression (e.g. **Supp. Fig. 4B**), we instead altered its specificity through mutations designed to disrupt B55 interactions with the downstream basic patch in its substrates (Cundell et al., 2016; Xu et al., 2008) (**Fig. 5C**). We generated mutations in complementary acidic residues in B55, changing these to alanine or creating charge swap substitutions of these sites to lysine. Compared to wild-type B55, ectopic expression of either of these mutants had a potent dominant effect on meiotic progression. Oocytes expressing the alanine mutant (DE/A) successfully assembled the meiosis I spindle and completed cytokinesis of polar body I. However, the meiosis II spindles did not form properly, displaying a ball-like morphology. Furthermore, anaphase and cytokinesis did not progress normally, resulting a reduced rate of successful polar body II extrusion (**Fig. 5D,E**). Oocytes expressing the charge-swap mutations (DE/K) displayed a more severe phenotype: the first meiotic spindle formed normally, but remained arrested, with the bivalent chromosomes maintaining cohesion and the co-oriented centromeres remaining fused, even at time points where wild-type control oocytes completed meiosis II. However, although normal anaphase I and cytokinesis did not occur, the spindle poles eventually separated, ultimately resulting in two semi-distinct spindles. Staining for Centrin2 revealed that pole fragmentation was due to the separation of the pair of centrioles in the spindle pole (**Fig. 5E**). Therefore, alterations to PP2A-B55 substrate specificity allow meiosis I events to occur, but prevent changes to the cell division machinery that occur at the MI/MII transition. We therefore propose that PP2A-B55 serves as essential regulator of the MI/MII transition by selectively dephosphorylating substrates to achieve exit from meiosis I, but retaining sites that must remain phosphorylated for meiosis II to occur.

### Threonine-specific dephosphorylation is essential for the spatiotemporal control of PP2A-B55 substrates

The selective TP dephosphorylation at the MI/MII transition suggests a potential mechanism to encode MI or MII-specific functions directly into substrates. Evolving phosphorylation sites as higher or lower affinity substrates for PP2A-B55 may provide temporal control for individual protein behaviors in meiosis. We identified several conserved phosphorylation sites on PRC1 (T470 in humans, T411 in sea star), TPX2 (T369 in humans, T508 in sea star), and INCENP (T59 in humans, T61 in sea star) (**Supp. Fig. 5B-D**) that are known substrates of PP2A-B55 (Cundell et al., 2016; Hein et al., 2017; Hummer and Mayer, 2009; Jiang et al., 1998). Consistent with the reactivation of PP2A-B55, these substrates display a decrease in phosphorylation following MI. As proof of principle for this model, we focused on the chromosome passenger complex (CPC) subunit INCENP, which contains both stable serine phosphorylations as well as MI-specific threonine phosphorylations (**Fig. 6A**). During anaphase of mitosis, the CPC transitions from the inner centromere to the central spindle, where it is required for cytokinesis (Carmena et al., 2012; Kaitna et al., 2000). The relocalization of human INCENP is opposed by phosphorylation on T59 (T61 in sea star), and requires dephosphorylation of this residue by PP2A-B55 (**Supp. Fig. 5D**) (Goto et al., 2006; Hummer and Mayer, 2009). However, the localization dynamics and impact of phosphorylation are not well defined in meiosis. We find that phosphorylation on T61 sharply decreased following MI, whereas other proline-directed sites on INCENP remained phosphorylated throughout the oocyte-to-embryo transition (**Fig. 6A**). Sequence alignment of sea star T61 with human INCENP T59 revealed the presence of a conserved downstream basic patch, supporting its potentially conserved dephosphorylation by PP2A-B55 (**Fig. 6B**).

**Figure 6.**
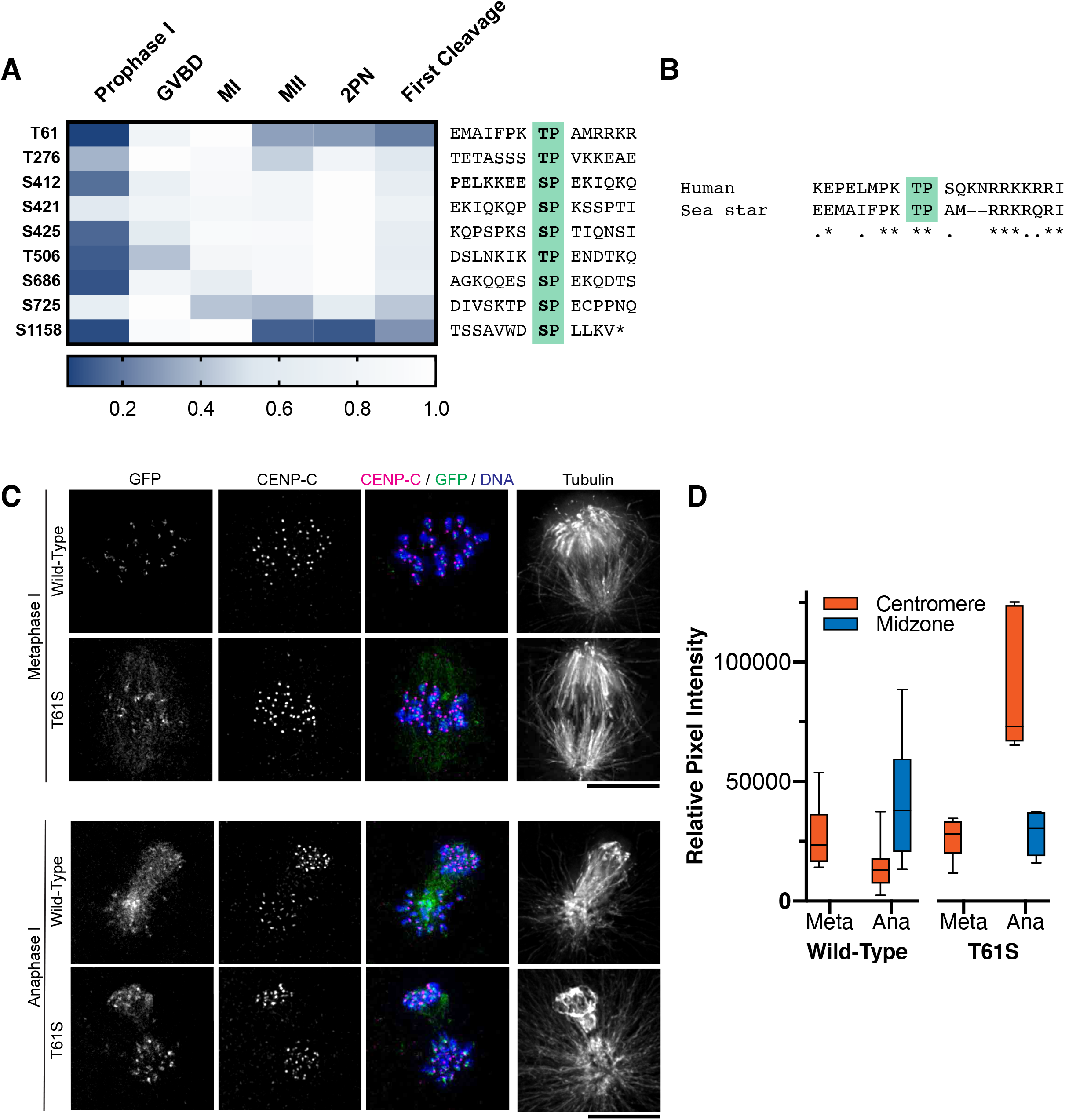
Dephosphorylation of a conserved threonine is necessary for INCENP localization. (A) Heatmap representation of relative phosphorylation levels of sites identified by phosphoproteomics. Phospho-centered amino acid sequences are provided on the right. (B) Sequence alignment of sea star T61 with human T59. (C) Immunofluorescence images of oocytes expressing wild-type or T61S mutant INCENP-GFP in Meiosis I. Wild-type INCENP translocates from centromeres to the central spindle in anaphase, whereas INCENP^T61S^ increases at centromere localization but not at the central spindle. Images are scaled individually to aid in visualization of localization changes. (D) Pixel intensity quantitation of unscaled oocyte images at centromeres or central spindles in metaphase (Meta) or anaphase (Ana) of meiosis I (WT meta n=8, WT ana n=8, T61S meta n=14, T61S ana n=6 oocytes).

To test the role of this INCENP threonine residue in meiosis I, we used GFP fusion constructs with either wild-type PmINCENP, or with T61 replaced with serine (T61S), which is typically considered to be a conservative change that would preserve protein function. Wild-type INCENP localized to centromeres in metaphase of meiosis I, but then relocalized to the central spindle at anaphase of Meiosis I. In contrast, INCENP(T61S)-GFP failed to translocate to the central spindle in anaphase of MI and remained at high levels at centromeres (Fig. 6C,D). The retention of INCENP at centromeres in anaphase I suggests that this residue remains phosphorylated at the MI/MII transition when substituted with serine, a lower affinity substrate for PP2A-B55. Thus, the usage of serine versus threonine, modulated by adjacent charged amino acids, can directly encode substrates with differential responses to a common set of kinases and phosphatases, enabling rewiring events at key cell cycle transitions. Through this paradigm, the behaviors of individual proteins may be temporally coordinated with meiotic cell cycle progression to achieve specific behaviors.

## Discussion

In this work, we define an extensive program of phosphorylation changes during the oocyte-to-embryo transition, spanning the complete developmental window from Prophase I arrest to the first embryonic cleavage. Using TMT-based proteomics, phosphoproteomics, and functional approaches, we find that overall protein levels are stable, but that selected cell cycle regulators, including Cyclins A and B, must undergo new protein synthesis for progression through meiosis. In addition, we identify a complex landscape of phosphorylation and dephosphorylation that underlies this developmental period. We find that the Prophase I arrest is characterized by low overall phosphorylation, and that maintaining this arrest requires phosphatase activity. Strikingly, although serine and threonine residues are often considered as conserved and interchangeable, we observe distinct behaviors for TP and SP phosphorylation at the MI/MII transition, with threonine sites being preferentially dephosphorylated. Such differential dephosphorylation suggests a novel paradigm for the regulatory control of oocytes, which must rapidly transition between the two meiotic divisions for the specialized meiotic cell cycle.

Our analysis reveals that upon decrease in Cdk activity at the MI/MII transition, selective dephosphorylation of TP versus SP sites by PP2A-B55 is essential for meiotic progression. Examination of the TP sites dephosphorylated after MI identified an enrichment for basic amino acids starting in the +2 position, matching the known consensus for PP2A-B55 substrates (Cundell et al., 2016; Kruse et al., 2020). Moreover, we find that PP2A-B55 is re-activated at the MI/MII transition, based on phosphorylation signatures of its inhibitory pathway, including Greatwall and ARPP19, as well as conserved PP2A-B55 substrates. Our results are consistent with prior work reporting a decrease in Cyclin B abundance and dephosphorylation of Greatwall kinase following MI and during first cleavage (Hara et al., 2012). Thus, the temporal profile of PP2A-B55 activity leaves it poised to play an essential role in the MI/MII transition for specializing the meiotic cell cycle and division machinery.

Our work supports the emerging picture that threonine and serine phosphorylation is not interchangeable (Cundell et al., 2016; Deana et al., 1982; Deana and Pinna, 1988; Hein et al., 2017; Pinna et al., 1976), but instead represent an important regulatory mechanism to temporally control cellular processes. In cultured mitotic cells, threonine dephosphorylation is important for timely mitotic exit. For example, inhibitory phosphorylations on the APC/C regulator Cdc20 are conserved as threonines, whereas serine substitution mutants display delayed dephosphorylation and delayed Cdc20 activation (Hein et al., 2017). Our work shows that this differential dephosphorylation can provide not just a kinetic delay, but also can confer distinct behaviors for the two different meiotic phases. For example, in our mutant analysis, serine substitution of threonine 61 for serine on INCENP, which is subject to inhibitory phosphorylation by Cdk1, resulted in its failure to translocate to the central spindle in anaphase of MI. These findings reveal the importance for temporally-coordinated dephosphorylation during meiotic progression, such that critical events including chromosome segregation and cytokinesis are properly synchronized with other cellular events. Such differential dephosphorylation represents a novel paradigm for the regulatory control of oocytes, which must rapidly transition between the two meiotic divisions for the specialized meiotic cell cycle.

These results point to PP2A-B55 as a critical conductor of cell cycle progression in oocytes. In diverse species, ARPP19/ENSA is required for exit from Prophase I arrest by inhibiting PP2A-B55 (Dupre et al., 2013; Dupré et al., 2014; Matthews and Evans, 2014; Okumura et al., 2014; Von Stetina et al., 2008). Our data in the sea star indicate that phosphatase activity is critical not only for maintaining the Prophase I arrest, but also to drive the transition between the two meiotic divisions. CDK and MAPK, combined with a wave of opposing PP2A-B55 activity at the MI/MII transition provides a pacemaker for meiotic cell cycle progression. By encoding PP2A-B55 affinity into substrates, different temporal phosphorylation profiles can be achieved, yielding subsets of cell division proteins that are constitutively active, or others that are uniquely modified in MI or MII. The rapidity between the two meiotic divisions (only 30 minutes in sea stars) places a selective pressure for specific substrates to be promptly dephosphorylated. In contrast, other sites may need to remain phosphorylated to prevent exit into interphase, creating evolutionary pressure for conservation as serine. The ability to segregate heritable material through meiosis is essential for organismal fitness. We propose that evolution has fine-tuned substrate dephosphorylation by selecting for amino acid sequences that favor or disfavor the interaction with PP2A-B55, thereby enabling precise temporal coordination of events in the oocyte-to-embryo transition.

## Methods and Materials

### Experimental Model and Subject Details

Sea stars (*Patiria miniata*) were wild-caught by South Coast Bio Marine (http://scbiomarine.com/) and kept in artificial seawater aquariums at 15 °C. Intact ovary and testis fragments were surgically extracted as previously described (Swartz et al., 2019).

### Ovary and Oocyte Culture

Ovary fragments were maintained in in artificial seawater containing 100 units/mL pen/strep solution. Intact ovary fragments were cultured this way for up to 1 week until oocytes were needed, with media changes every 2-3 days. Isolated oocytes were cultured for a maximum of 24 hours in artificial seawater with pen/strep. To induce meiotic reentry, 1-methyladenine (Acros Organics) was added to the culture at a final concentration of 10 μM. For fertilization, extracted sperm was added to the culture at a 1:1,000,000 dilution. For emetine treatments (**Fig. 2**), oocytes were pre-treated with 10 μM emetine (Sigma-Aldrich) for 30 minutes prior to hormonal stimulation.

### Mass Spectrometry Sample Preparation

Oocytes were collected by centrifugation at 200 x g and resuspending pellets and washed one time with wash buffer (50 mM HEPES, pH 7.4, 1 mM EGTA, 1 mM MgCl_2_, 100 mM KCl, 10% glycerol), pelleted with all excess buffer removed, and snap frozen in liquid nitrogen. Samples were lysed and proteins were digested into peptides with trypsin. Oocyte pellets were lysed in ice-cold lysis buffer (8 M urea, 25 mM Tris-HCl pH 8.6, 150 mM NaCl, phosphatase inhibitors (2.5 mM beta-glycerophosphate, 1 mM sodium fluoride, 1 mM sodium orthovanadate, 1 mM sodium molybdate) and protease inhibitors (1 mini-Complete EDTA-free tablet per 10 ml lysis buffer; Roche Life Sciences)) and sonicated three times for 15 sec each with intermittent cooling on ice. Lysates were centrifuged at 15,000 x g for 30 minutes at 4°C. Supernatants were transferred to a new tube and the protein concentration was determined using a BCA assay (Pierce/ThermoFisher Scientific). For reduction, DTT was added to the lysates to a final concentration of 5 mM and incubated for 30 min at 55°C. Afterwards, lysates were cooled to room temperate and alkylated with 15 mM iodoacetamide at room temperature for 45 min. The alkylation was then quenched by the addition of an additional 5 mM DTT. After 6-fold dilution with 25 mM Tris-HCl pH 8, the samples were digested overnight at 37°C with 1:100 (w/w) trypsin (Promega). The next day, the digest was stopped by the addition of 0.25% TFA (final v/v), centrifuged at 3500 x g for 15 min at room temperature to pellet precipitated lipids, and peptides were desalted. Peptides were lyophilized and stored at −80°C until further use. For proteomics analysis, an aliquot containing ~ 100 μg of peptides were resuspended in 133 mM HEPES (SIGMA) pH 8.5, TMT reagent (Thermo Scientific) stored in dry acetonitrile (ACN) (Burdick & Jackson) was added, vortexed to mix reagent and peptides. After 1 hr at room temperature, an aliquot was withdrawn to check for labeling efficiency while the remaining reaction was stored at −80°C. Once labeling efficiency was confirmed to be at least 95%, each reaction was quenched by addition of ammonium bicarbonate to a final concentration of 50mM for 10 minutes, mixed, diluted with 0.1% TFA in water, and desalted. The desalted multiplex was dried by vacuum centrifugation and separated by offline Pentafluorophenyl (PFP)-based reversed phase HPLC fractionation as previously described (Grassetti et al., 2017).

For phosphoproteomic analysis, ~4 mg of peptides were enriched for phosphopeptides using a Fe-NTA phosphopeptide enrichment kit (Thermo Scientific) according to instructions provided by the manufacture and desalted. Phosphopeptides were labeled with TMT reagent and labeling efficiency was determined as described above. Once labeling efficiency was confirmed to be at least 95%, each reaction was quenched, mixed, diluted with 0.1% TFA in water, and desalted. The desalted multiplex was dried by vacuum centrifugation and separated by offline Pentafluorophenyl (PFP)-based reversed phase HPLC fractionation as previously described (Grassetti et al., 2017).

Proteomic and phosphoproteomic analyses of oocyte to embryo time courses was performed on an Orbitrap Fusion mass spectrometer (Thermo Scientific) equipped with an Easy-nLC 1000 (Thermo Scientific) (Senko et al., 2013). Peptides were resuspended in 8% methanol / 1% formic acid across a column (40 cm length, 100 μm inner diameter, ReproSil, C18 AQ 1.8 μm 120 Å pore) pulled in-house across a 2 hr gradient from 3% acetonitrile/0.0625% formic acid to 37% acetonitrile/0.0625% formic acid. The Orbitrap Fusion was operated in data-dependent, SPS-MS3 quantification mode (McAlister et al., 2014; Ting et al., 2011)For proteomics analysis, an Orbitrap MS1 scan was taken (scan range = 350 – 1300 m/z, R = 120K, AGC target = 2.5e5, max ion injection time = 100ms). Followed by data-dependent iontrap trap MS2 scans on the most abundant precursors for 3 seconds in rapid scan modem, AGC target = 1e4, max ion injection time = 40ms, CID collision energy = 32%). Followed by an Orbitrap MS3 scans for quantification (R = 50K, AGC target = 5e4, max ion injection time = 100ms, HCD collision energy = 60%, scan range = 110 – 750 m/z, synchronous precursors selected = 10). For phosphoproteomics analysis, an Orbitrap MS1 scan was taken (scan range = 350 – 1300 m/z, R = 120K, AGC target = 35e5, max ion injection time = 100ms). Followed by data-dependent Orbitrap trap MS2 scans on the most abundant precursors for 3 seconds Ion selection; charge state = 2: minimum intensity 2e5, precursor selection range 625-1200 m/z; charge state 3: minimum intensity 3e5, precursor selection range 525-1200 m/z; charge state 4 and 5: minimum intensity 5e5). Quadrupole isolation = 0.7 m/z, R = 30K, AGC target = 5e4, max ion injection time = 80ms, CID collision energy = 32%). Followed by an Orbitrap MS3 scans for quantification (R = 50K, AGC target = 5e4, max ion injection time = 100ms, HCD collision energy = 65%, scan range = 110 – 750 m/z, synchronous precursors selected = 5).

The proteomic analysis of emetine treated oocytes was performed on an Orbitrap Lumos mass spectrometer (Thermo Scientific) equipped with an Easy-nLC 1200 (Thermo Scientific). Peptides were resuspended in 8% methanol / 1% formic acid across a column (40 cm length, 100 μm inner diameter, ReproSil, C18 AQ 1.8 μm 120 Å pore) pulled inhouse across a 2 hr gradient from 3% acetonitrile/0.0625% formic acid to 37% acetonitrile/0.0625% formic acid. The Orbitrap Lumos was operated in data-dependent, SPS-MS3 quantification mode (McAlister et al., 2014; Ting et al., 2011). An Orbitrap MS1 scan was taken (scan range = 350 – 1400 m/z, R = 120K, AGC target = 2.5e5, max ion injection time = 50ms), followed by data-dependent ion trap MS2 scans on the most abundant precursors for 1.8 seconds. Ion selection; charge state = 2: minimum intensity 2e5, precursor selection range 600-1400 m/z; charge state 3-5: minimum intensity 4e5). Quadrupole isolation = 0.8 m/z, CID collision energy = 35%, CID activation time = 10 ms, activation Q = 0.25, scan range mode = m/z normal, ion trap scan rate = rapid, AGC target = 4e3, max ion injection time = 40 ms). Orbitrap MS3 scans for quantification (R = 50K, AGC target = 5e4, max ion injection time = 100ms, HCD collision energy = 65%, scan range = 100 – 500 m/z, synchronous precursors selected = 10).

The phosphoproteomic analysis of Calyculin A treated occytes was performed on the Orbitrap Lumos mass spectrometer. Here, wherein an Orbitrap MS1 scan was taken (scan range = 350 – 1250 m/z, R = 120K, AGC target = 2.5e5, max ion injection time = 50ms), followed by data-dependent Orbitrap MS2 scans on the most abundant precursors for 2 seconds. Ion selection; charge state = 2: minimum intensity 2e5, precursor selection range 650-1250 m/z; charge state 3: minimum intensity 3e5, precursor selection range 525-1250 m/z; charge state 4 and 5: minimum intensity 5e5). Quadrupole isolation = 1 m/z, R = 30K, AGC target = 5e4, max ion injection time = 55ms, CID collision energy = 35%). Orbitrap MS3 scans for quantification (R = 50K, AGC target = 5e4, max ion injection time = 100ms, HCD collision energy = 65%, scan range = 100 – 500 m/z, synchronous precursors selected = 5).

The raw data files were searched using COMET with a static mass of 229.162932 on peptide N-termini and lysines and 57.02146 Da on cysteines, and a variable mass of 15.99491 Da on methionines against a target-decoy version of the *Patiria miniata* proteome sequence database. For phosphoproteomics analysis 79.96633 Da on serines, threonines and tyrosines was included as an additional variable mass. Data and filtered to a <1% FDR at the peptide level. Quantification of LC-MS/MS spectra was performed using in house developed software. Probability of phosphorylation site localization was determined by PhosphoRS (Taus et al., 2011).

The *Patiria miniata* proteome sequence database was generated using published paired-end RNA-seq data for a Patiria miniata ovary sample (SRX445851 / SRR1138705), with the Agalma transcriptome pipeline (Dunn et al., 2013). Each protein sequence from this transcriptome was assessed for similarity with known proteins in the NCBI nr (nonredundant) sequence database using NCBI BLAST. Transcriptome sequences were annotated with the accession and sequence similarity metrics of their top BLAST hits. This annotation procedure was repeated for an unannotated protein database sourced from EchinoBase to generate a composite database. Duplicate sequences and subsequences within the composite database were removed to reduce redundancy.

### Construct Generation

*Patiria miniata* INCENP was identified using the genomic resources at echinobase.org and previously published ovary transcriptomes (Kudtarkar and Cameron, 2017; Reich et al., 2015). Wild-type INCENP was amplified from first strand cDNA reverse transcribed from total ovary mRNA. T61A, S, and D mutants were generated by overlap extension PCR. These cDNAs were then cloned into pCS2+8 as c-terminal GFP fusions (Gokirmak et al., 2012). Wild-type and mutant versions of B55 were synthesized (Twist Biosciences) and cloned into pCS2+8 with standard restriction enzyme methods.

### Oocyte Microinjection

For expression of constructs in oocytes, plasmids were linearized with NotI to yield linear template DNA. mRNA was transcribed in vitro using mMessage mMachine SP6 and the polyadenylation kit (Life Technologies), then precipitated using lithium chloride solution. Prophase I arrested oocytes were injected horizontally in Kiehart chambers with approximately 10-20 picoliters of mRNA solution in nuclease free water. B55 constructs were injected at 500 ng/μl, and INCENP constructs were injected at 1000 ng/μl. After microinjection, oocytes were cultured 18-24 to allow time for the constructs to translate before 1-methyladenine stimulation. Custom synthesized cyclin morpholinos, or the Gene Tools standard control, were injected at 500 μM immediately before 1-methyladenine stimulation (Gene Tools).

### Immunofluorescence, Imaging, and Immunoblots

Oocytes were fixed at various stages in a microtubule stabilization buffer as described previously (2% paraformaldehyde, 0.1% Triton X-100, 100 mM HEPES, pH 7.0, 50 mM EGTA, 10 mM MgSO_4_, 400 mM dextrose) for 24 hours at 4° C (von Dassow et al., 2009). The oocytes were then washed 3 times in PBS with 0.1% Triton X-100, and blocked for 15 minutes in AbDil (3% BSA, 1 X TBS, 0.1% triton X-100, 0.1% Sodium Azide). Primary antibodies diluted in AbDil were then applied and the oocytes were incubated overnight at 4° C. Anti-CENP-C and alpha tubulin (DM1α, Sigma) antibodies were used at 1:5,000. DNA was stained with Hoechst. GFP booster (Chromotek) was used at 1:500 to improve the signal from GFP fusion constructs. The oocytes were compressed under coverslips in ProLong Gold antifade Mountant (Thermo Fisher). Images were collected with a DeltaVision Core microscope (Applied Precision/GE Healthsciences) with a CoolSnap HQ2 CCD camera and a 100x 1.40 NA Olympus U-PlanApo objective. Confocal images (**Fig. 2G**) were collected with a Yokogawa W1 spinning disk microscope. Images were processed with Fiji, and scaled equivalently across conditions unless otherwise specified (Schindelin et al., 2012). For Western blot analyses, oocytes were lysed in Laemmli samples buffer, separated by SDS-PAGE, and analyzed by western blotting. Antibodies were diluted in 3% milk in TBST and phosphatase inhibitors. Anti phospho Cdk1 Y15 (Cell Signaling #4539), anti phospho ERK1/2 T202/Y204 (Cell Signaling #9101, this antibody detect singly and dually phosphorylated ERK1/2 and the total intensity of singly and dually phosphorylated ERK1/2 in sea start suggest that Y193 is the dominant phosphoform), and phospho CDK Substrate [pTPXK] (Cell Signaling, #14371). Antibodies raised against phosphorylation sites on human proteins were evaluated for cross-reactivity and judged to do if they 1) only recognized one band, 2) the size of the band migrated at the expected molecular weight, and 3) the phosphorylation pattered matched the MS quantification.

### Quantification and Statistical Analysis

For the time course proteomic analysis, peptide TMT intensities were summed on a per protein basis. Replicate multiplexes were normalized to each other based on the internal bridge sample. Protein intensities were adjusted based on total TMT reporter ion intensity in each channel. Individual protein intensity across each time course were scaled between 0 and 1. For the time course phosphoproteomic analysis, replicate multiplexes were normalized to each other based on the internal bridge sample. Phosphopeptide intensities were adjusted based on total amount of protein input and median total TMT reporter ion intensity in each channel per condition. Individual phosphopeptide intensity across each time course were scaled between 0 and 1. A Pearson correlation coefficient for the time course data was calculated as pairwise correlation for proteins and phosphopeptides identified in two time course series and as multiple correlations for proteins identified in three time course series in Excel. A P-value was calculated from the Pearson correlation coefficient. Proteins quantified in all three time course series were analyzed by hierarchical clustering using Euclidean distance and average linkage in Perseus. Phosphorylation sites with a localization score of 0.9 or higher and quantified with a correlation coefficient of 0.8 more (corresponding to a p-value of 0.05 or less) were analyzed by hierarchical clustering using Euclidean distance and average linkage in Perseus. For the Calyculin A treatment, the 0 min and 30 min and the 0 min and 70 min time points were compared. P-values were calculated using a two-tailed Student’s T-test assuming unequal variance. Average phosphorylation abundances of phosphorylation site upon 0, 30, and 70 min Calyculin A treatment were compared between Prophase 1 and other phases by 2×3 Fisher Exact test. For the emetine proteomics analysis, control and emetine-treated oocytes were compared at the MI and 2-PN stage. Peptide TMT intensities were summed on a per protein basis. Protein intensities were adjusted based on total TMT reporter ion intensity in each channel and log-transformed. P-values were calculated using a two-tailed Student’s T-test assuming unequal variance.

For gene ontology analysis for proteins significantly changes in abundance during oocyte to embryo transition or emetine treatment, human homologs of *Patiria miniata* proteins were identified using NCBI’s BLAST+. A custom R script was then used to filter the BLAST results to only high-confidence matches (E-value < .01) and to remove redundant matches. The resulting list analyzed using WebGestaltR to find enriched GO terms (Liao et al., 2019). Motif analysis for selected and deselected amino acids surrounding a phosphorylation site was performed using the Icelogo web interface. Statistical analysis to determine population difference of dephosphorylation preferences at the MI to MII transition was performed using an unpaired nonparametric Kolmogorov-Smirnov (KS) T test.

Graphing and statistical analyses involving the scoring of meiotic phenotypes (e.g. Fig. 5D) were performed using Prism (GraphPad). The statistical tests used to calculate p-values are indicated in the figure legends. Pixel intensity quantifications of INCENP fluorescence images (i.e. Fig. 6D) were conducted using Fiji. Circular ROIs were placed over inner centromeres or the central spindle (7px and 30px diameter, respectively), along with an adjacent ROI to measure the background intensity of equivalent size by RawIntDen. The background measurements were then subtracted from the centromere or midzone measurement, and the resulting value was normalized to the selection area to allow comparison between inner centromere and midzone intensity. The crystal structure of the PP2A-B55 holoenzyme (**Fig. 5A**) was displayed using UCSF ChimeraX (Goddard et al., 2018).

## Data and materials availability

Raw MS data for the experiments performed in this study are available at MassIVE and ProteomeXchange. Plasmids generated from this study are available on request and will be deposited to Addgene.

## Acknowledgments

S.Z.S. was supported by a K99 fellowship from NIH/NICHD (5K99HD099315). This work was supported by grants from the NIH/National Institute of General Medical Sciences to I.M.C. (R35GM126930) and A.N.K (R35GM119455). The Orbitrap Fusion Tribrid mass spectrometer was acquired with support from NIH (S10-OD016212). Molecular graphics and analyses performed with UCSF ChimeraX, developed by the Resource for Biocomputing, Visualization, and Informatics at the University of California, San Francisco, with support from National Institutes of Health R01-GM129325 and the Office of Cyber Infrastructure and Computational Biology, National Institute of Allergy and Infectious Diseases.

## Supplemental Figure Legends

**Figure S1. Mass spectrometry samples and dataset statistics.** (A) Representative immunofluorescence images of oocytes at stages collected for mass spectrometry. Scale bars = 10μm. (B,C) Venn diagram representation of the proteins and phosphorylation events identified in three biological replicates, respectively. (D) Distribution of phosphorylation sites identified by phosophoproteomics on serine, threonine, and tyrosine.

**Figure S2. Mass spectrometry analysis of nascent protein synthesis.** (A,B) Volcano plots of protein abundance changes in emetine-treated oocytes versus controls when control oocytes were in meiosis II (A) or the pronuclear stage (B). Proteins with abundance changes of greater than 2 fold change and a p-value of less than 0.05 are colored green (108 out of 7,609 proteins).

**Figure S3. Proteomic and biochemical analysis of phosphatases.** (A) Phosphorylation profile of Greatwall kinase at T204, within its activation segment. (B) Phosphorylation of the conserved inhibitory site T316 on PP1G. (C,D) Phosphorylation of conserved NIPP1 sites that disrupt its interaction with PP1. Light blue represents the standard deviation (E) Sequence alignment of sea star NIPP1 with orthologs in *Danio rerior* (drer), *Canis lupus* (clup), *Equus caballus* (ecab), *Homo sapiens* (hsap), *Monodelphis domestica* (mdom), *Macaca mulatta* (mmul), *Mus musculus* (mmus), *Ornithorhynchus anatinus* (oana), *Pan troglodytes* (ptro), *Gallus gallus* (ggal). (F) Immunoprecipitation of wild-type or phosphonull NIPP1 mutants tagged with FLAG, followed by Western blot to test PP1 association. Calyculin treatment was performed to increase phosphorylation on NIPP1 that decreases association with PP1 (lane 4), but phosphonull mutations (S197A, S199A) increases the association with PP1 (compare lanes 6 and 8 to lane 4). (G) Sequence alignment of amino acids surrounding the Greatwall phosphorylation site on human ARPP19 and ENSA and sea star ARPP19. (H) Western blot of Cdc25 and pENSA in control cell lysate (lane 1), or lysate to which thiophosphorylated (lane 2) PmArpp19^WT^ or PmArpp19^S106A^ was added. Thiophosphorylated Arpp19^WT^ results in a phosphorylation shift of the band for PP2A-B55 substrate Cdc25, indicating inhibition of the phosphatase. (I) Temporal phosphorylation profile of the conserved Greatwall phosphorylation on ARPP19 S106. Light blue shading represents standard deviation.

**Figure S4. Calyculin A treatment of sea star oocytes** (A) DIC images of oocytes treated with DMSO control or 1 μM calyculin. Dose responsivity curve of percentage of oocytes undergoing germinal vesicle breakdown (GVBD) in response to different concentrations of Calyculin. Errors bars represent the standard error of the mean. Scale bars = 100μm (B) Immunofluorescence images for tubulin, CENP-C, and phalloidin staining for actin following stimulation with 1-methyladenine or calyculin. Scale bars = 10μm. (C) Hierarchical clustering of phosphorylation events (localization score of 0.9 or higher, p-value of 0.05 or less, and stage specific peak phosphorylation) clustered by phosphosite (vertically) and meiotic stage (columns) as in Figure 3B with phosphorylation changes upon Calyculin A treatment added. (D) Comparison of average phosphorylation abundance changes upon Calyculin A treatment for 0 min, 30 min, or 70 min in a stage-specific manner. Statistical comparison was performed by Fisher Exact test (*p-value <0.05).

**Figure S5. Phospho-threonine phosphorylation site analyses** (A) Table depicted all significant differences in the population distribution as determined by KS statistics in Figure 4E. Significant (p-value: *<0.05, **<0.01, ***<0.001, **** <0.0001). (B-D) Relative abundances and sequence alignment of PP2A-B55-dependent phosphorylation sites on PRC1, TPX2, and INCENP with orthologs in *Danio rerior* (drer), *Canis lupus* (clup), *Equus caballus* (ecab), *Homo sapiens* (hsap), *Monodelphis domestica* (mdom), *Macaca mulatta* (mmul), *Mus musculus* (mmus), *Ornithorhynchus anatinus* (oana), *Pan troglodytes* (ptro), *Gallus gallus* (ggal), *Takifugu rubripes* (trub) *Rattus norvegicus* (rnor). Light blue shading represents standard deviation.

## Supplemental Tables

**Supplemental Table 1:** Protein abundances quantified across time course proteomics from prophase I arrest to the first embryonic cleavage

**Supplemental Table 2:** Results of gene ontology analysis of proteins with significant changes in abundances from prophase I arrest to the first embryonic cleavage

**Supplemental Table 3:** Protein abundances changes upon emetine treatment

**Supplemental Table 4:** Results of gene ontology analysis of proteins with significant changes upon emetine treatment

**Supplemental Table 5:** Phosphorylation abundances quantified across time course proteomics from prophase I arrest to the first embryonic cleavage and upon Calyculin A treatment

