## Supplemental Figures for "Selective dephosphorylation by PP2A-B55 directs the meiosis I - meiosis II transition in oocytes"

### Supp. Figure 1

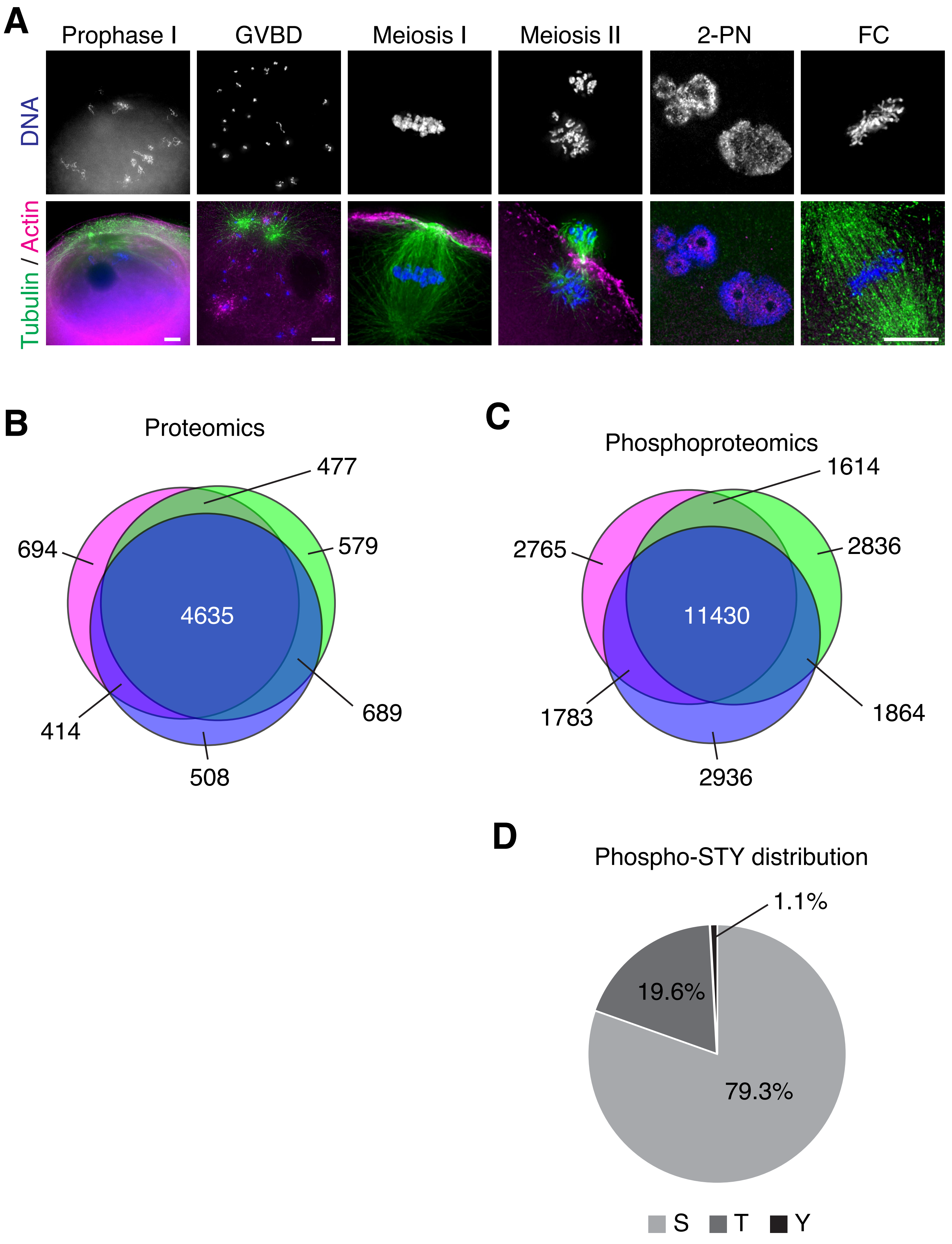

### Supp. Figure 2

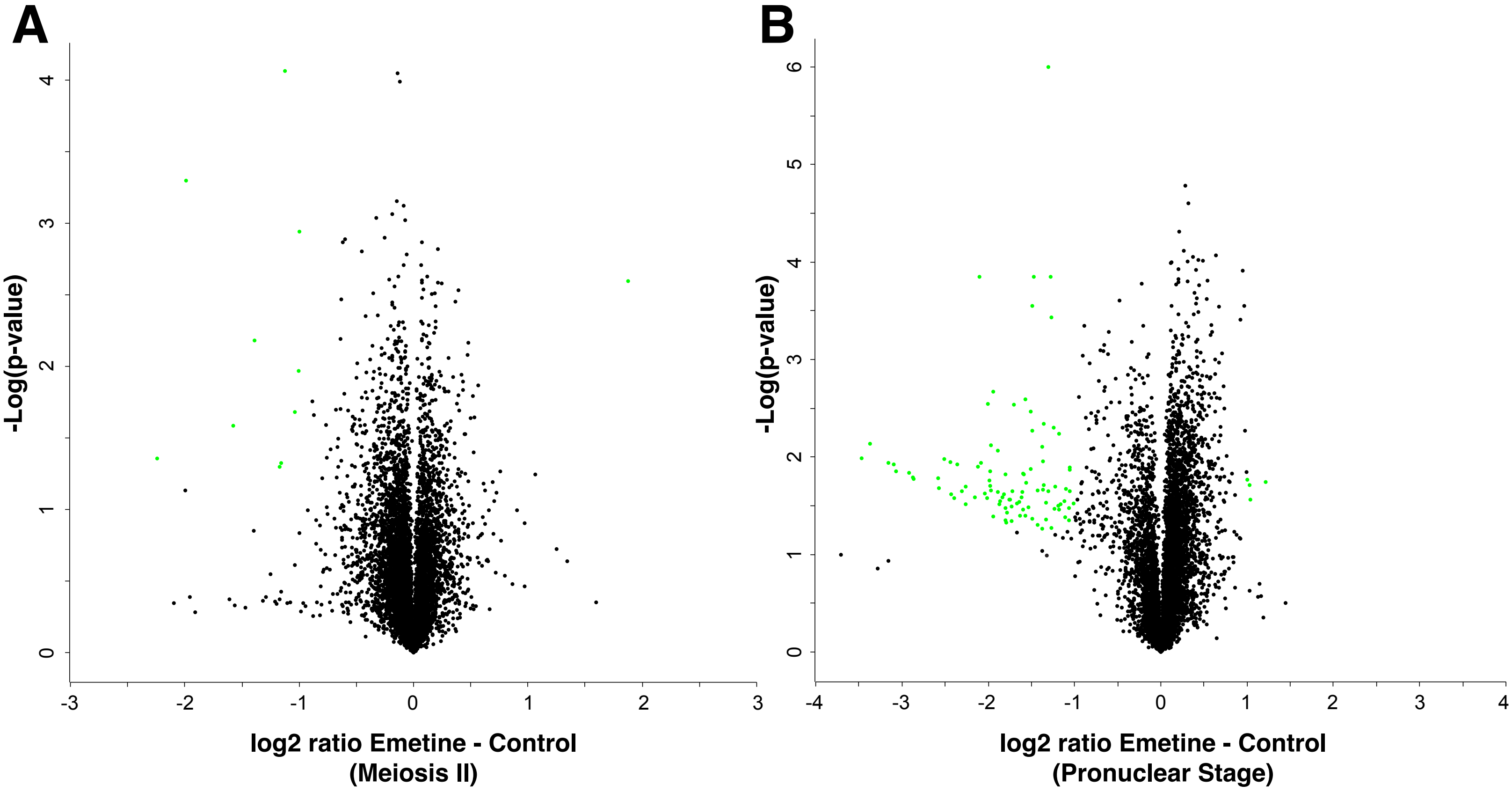

Supp. Figure 3

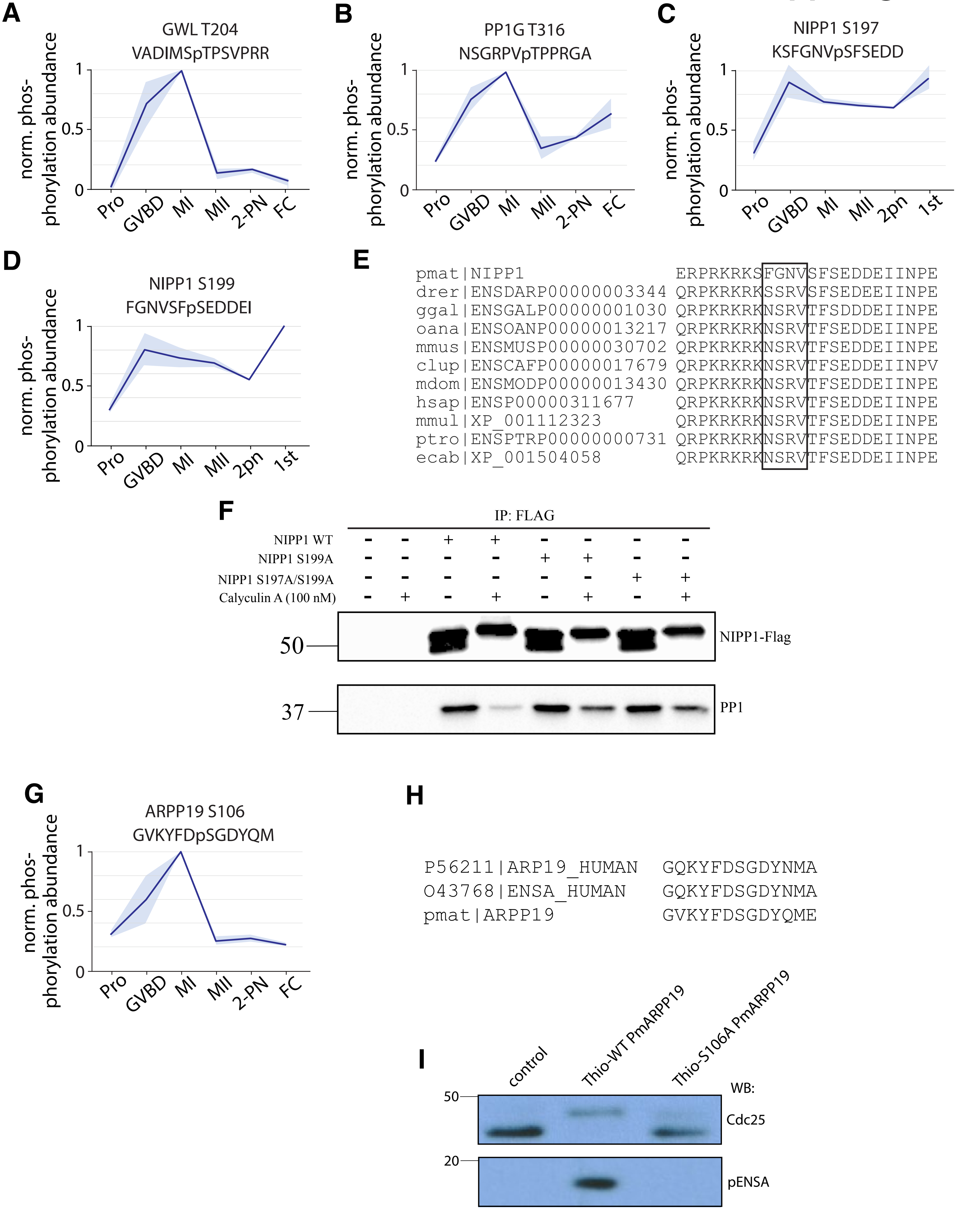

Supp. Figure 4

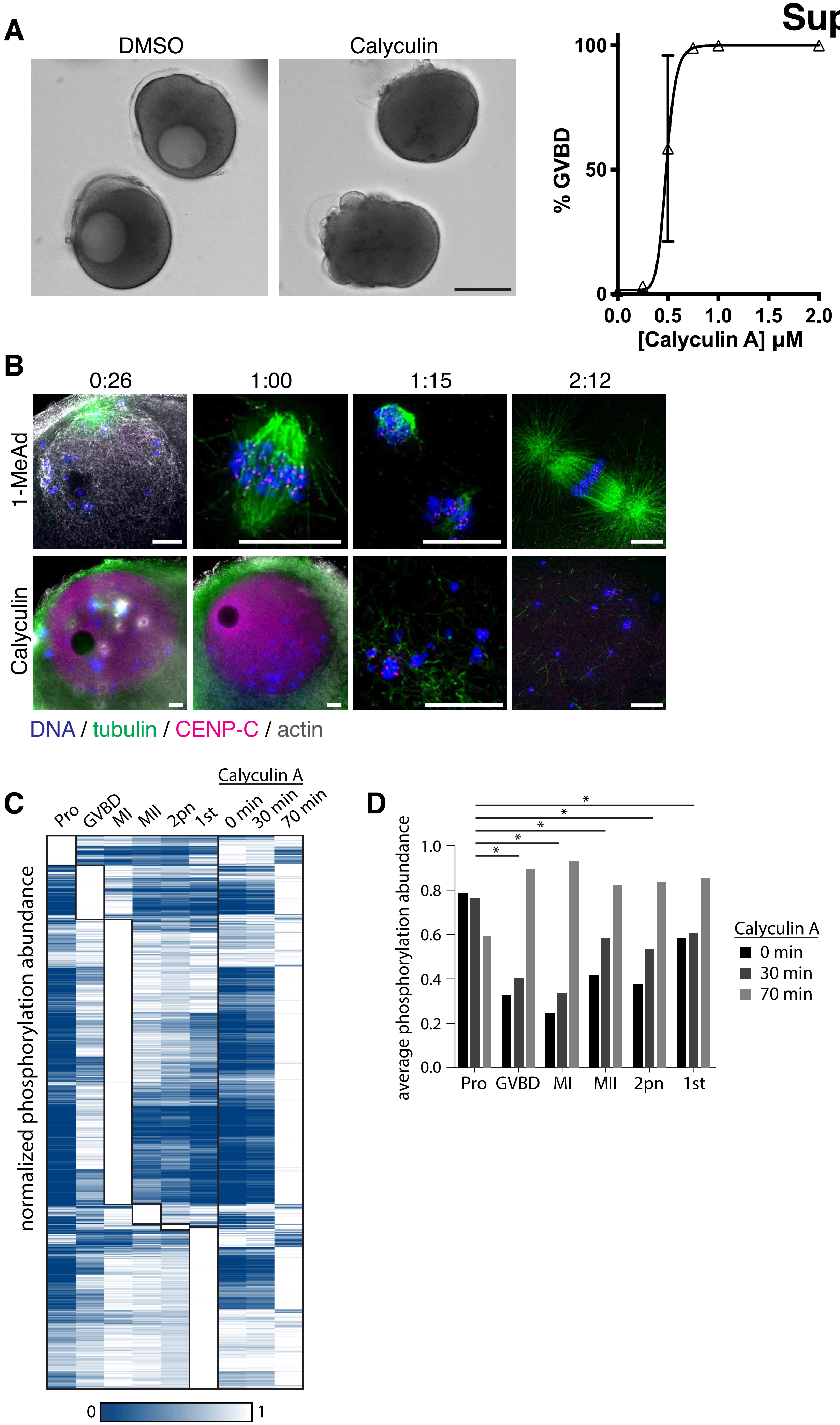

### Supp. Figure 5

**A**

| | TPA | TPD | TPE | TPG | TPK | TPP | TPR |
|---|---|---|---|---|---|---|---|
| TPA | | * | **** | | | **** | |
| TPD | * | | | | | | |
| TPE | **** | | | * | ** | | ** |
| TPG | | | * | | | *** | |
| TPK | | | ** | | | **** | |
| TPP | **** | | | *** | **** | | **** |
| TPR | | | ** | | | **** | |

**B**

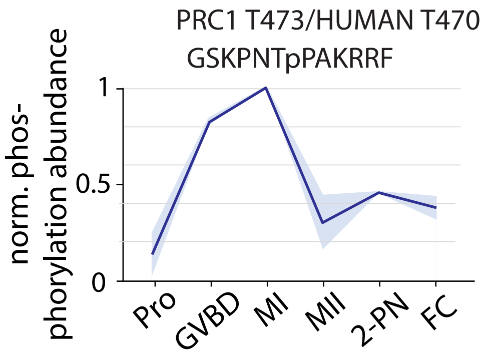


```
pmat|Pmi_640672              NEMTFGSKPNTPAKRRFNGNS
drer|ENSDARP00000089594      EDMVYGTVIRTPTKRRFLGST
ggal|ENSGALP00000028134      TEMMYGSTPRTPIKRRLLGPH
mdom|ENSMODP00000024319      TEMLYGSTPRTPGKRRGPGPN
mmus|ENSMUSP00000043379      AEMLYGSTPRTPSKRPGQTPK
ecab|XP_001498877            TEMLYGSAPRTPNKRRGLAPN
clup|ENSCAFP00000017901      TEMLYGSIPRTPNKRQGLTPN
hsap|ENSP00000377793PRC1     TEMLYGSAPRTPSKRRGLAPN
mmul|XP_001098763            TEMLYGSAPRTPSKRRGLAPN
```

**C**

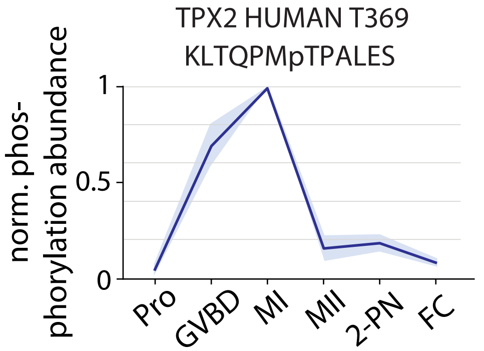


```
pmat|Pmi_661850              KKAKL-TQPMTPALESRNRHR
drer|ENSDARP00000101360      DKSKI-THPKTPQLLTQQRHR
trub|ENSTRUP00000023233      DHLKL-TQPHTPHLVTKQRSR
oana|ENSOANP00000020123      NAPKL-TQPRTPSFQTKGRLR
mdom|ENSMODP00000024312      SMLKI-TNPQTPVFKTKQRTR
mmus|ENSMUSP00000105441      SVNKIARDPQTPILQTKYRTR
rnor|ENSRNOP00000010851      PVNKIVRDPQTPILQTKYRAR
mmul|XP_001109833            SVTKICRDPQTPVLQTRHRAR
hsap|ENSP00000341145         SVTKICRDPQTPVLQTKHRAR
ptro|ENSPTRP00000022905      SVTKICRDPQTPVLQTKHRAR
clup|ENSCAFP00000010516      AGLKISRDPQTPVLQTKQRTR
ecab|XP_001499807            TVAKISRDPQTPVLQTKQRAR
```

**D**

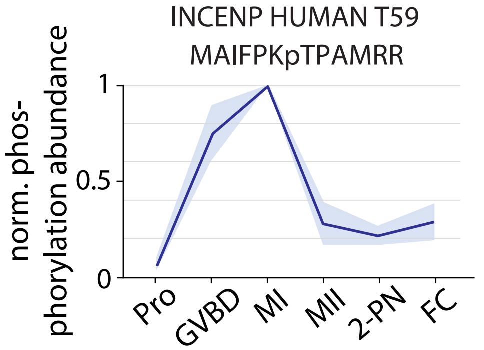


```
pmat|Pmi_669602              --EEMAIFPKTPAMRRKRQRI
trub|ENSTRUP00000019900      FNTEPELMPKTPSQK-KNSRR
ggal|ENSGALP00000012165      FRAEPGLMPKTPSQR-RRPRK
mdom|ENSMODP00000033394      FSVEPELMPKTPSQK-NRQKK
mmus|ENSMUSP00000108793      FSNEPELMPKTPSQK-NRRKK
rnor|ENSRNOP00000040392      FNNEPELMPKTPSQK-NRRKK
clup|ENSCAFP00000023376      FSKEPELMPKTPSQK-NRQKK
ecab|XP_001916204            FSKEPELMPKTPSQK-NRRKK
hsap|ENSP00000378295         FSKEPELMPKTPSQK-NRRKK
ptro|ENSPTRP00000059623      FSKEPELMPKTPSQK-NRRKK
mmul|XP_001118481            FSKEPELMPKTPSQK-NRRKK
```
